# Network-Based Matching of Patients and Targeted Therapies for Precision Oncology^*^

**DOI:** 10.1101/727941

**Authors:** Qingzhi Liu, Min Jin Ha, Rupam Bhattacharyya, Lana Garmire, Veerabhadran Baladandayuthapani

## Abstract

The extensive acquisition of high-throughput molecular profiling data across model systems (human tumors and cancer cell lines) and drug sensitivity data, makes precision oncology possible – allowing clinicians to match the right drug to the right patient. Current supervised models for drug sensitivity prediction, often use cell lines as exemplars of patient tumors and for model training. However, these models are limited in their ability to accurately predict drug sensitivity of individual cancer patients to a large set of drugs, given the paucity of patient drug sensitivity data used for testing and high variability across different drugs. To address these challenges, we developed a multilayer network-based approach to impute individual patients’ responses to a large set of drugs. This approach considers the triplet of patients, cell lines and drugs as one inter-connected holistic system. We first use the omics profiles to construct a patient-cell line network and determine best matching cell lines for patient tumors based on robust measures of network similarity. Subsequently, these results are used to impute the “missing link” between each individual patient and each drug, called <u>Personalized Imputed Drug Sensitivity Score</u> (PIDS-Score), which can be construed as a measure of the therapeutic potential of a drug or therapy. We applied our method to two subtypes of lung cancer patients, matched these patients with cancer cell lines derived from 19 tissue types based on their functional proteomics profiles, and computed their PIDS-Scores to 251 drugs and experimental compounds. We identified the best representative cell lines that conserve lung cancer biology and molecular targets. The PIDS-Score based top sensitive drugs for the entire patient cohort as well as individual patients are highly related to lung cancer in terms of their targets, and their PIDS-Scores are significantly associated with patient clinical outcomes. These findings provide evidence that our method is useful to narrow the scope of possible effective patient-drug matchings for implementing evidence-based personalized medicine strategies.

**Data and code availability:** https://github.com/bayesrx/bayesrx.github.io/tree/master/authors/liu-q./ Shiny app (data and results visualization tool): https://qingzliu.shinyapps.io/psb-app/

## 1. Introduction

Owing to the inherent tumor complexity and molecular heterogeneity, the main goal of precision oncology (or targeted cancer therapy) is to treat a patient by selectively inhibiting molecular (e.g., gene and protein) alterations that contribute to cancer development and progression.^1^ Recent advances in high-throughput multi-omic technologies (e.g. genomics, transcriptomics, proteomics and metabolomics) have accelerated the implementation of precision medicine-based strategies.^2^ Large-scale investigations of molecular, mutational and oncogenic landscapes across human cancers, such as The Cancer Genome Atlas (TCGA)^3^ and International Cancer Genome Consortium (ICGC)^4^ have provided extensive molecular characterization of thousands of cancer and normal samples spanning >30 cancer types. These datasets have a rich catalogue of molecular alterations that provide an opportunity to quantify and assess drug responses in patients within and across tumor types. On the other hand, since the clinical datasets of patient drug responses are time-consuming to generate, expensive and limited in the scope of drugs that can be tested on patients, these datasets are still relatively small and present challenges to be used as reliable training data for drug sensitivity prediction models.^5^ As an alternative, researchers have sought to match patients to effective drug therapies by generating large pharmacogenomics databases from tumor-derived cell lines, such as Cancer Cell Line Encyclopedia (CCLE)^6^ and Genomics of Drug Sensitivity in Cancer (GDSC)^7^, based on the premise that cell lines can serve as suitable *in-vitro* models to understand the therapeutic responses of patient tumors to drugs.^8^

Recently a few studies have used molecular profiles of cancer cell lines to fit supervised machine learning models and predict drug responses in patients.^9–10^ However, since the drug sensitivity data used for testing are available for only a few drugs and the model performance has high variability across different drugs, these models are limited in their ability to accurately predict the drug sensitivity of individual cancer patients over a large set of drugs. In addition, it would be time-consuming and costly to conduct preclinical (or experimental) tests to validate all possible patient-drug combinations. These challenges underscore the urgent need for developing reliable *in-silico* methods to examine or re-evaluate the drug responses of individual cancer patients to a large set of drugs, in order to narrow down the number of optimal therapies for implementing evidence-based personalized medicine strategies.

To address the above challenges, we developed a multilayer network-based approach to impute individual patients’ responses to a large set of drugs, by considering the triplet of patients, cell lines and drugs as one inter-connected holistic system. In the conceptual scheme shown in Figure 1(a), cell lines serve as an intermediary layer to assess sensitivities of drugs (bottom layer) in their highly matching patients (top layer). The analysis consists of two main steps as shown in Figure 1(b). First, patients and cell lines are matched based on a multiscale community detection algorithm^11^, which investigates topological structures of the patient-cell line network at multiple scales. This allows for a more efficient assimilation of network-connectivity information to obtain matching scores, which we call adjusted co-clustering scores, between patient tumors and cell lines. In the second step, a locally linear weighted model is used to impute the “missing link” between each individual patient and each drug, called <u>Personalized Imputed Drug Sensitivity Score</u> (PIDS-Score). This score can be construed as a measure of the therapeutic potential of a drug or therapy. To compute the PIDS-Score of a patient to a specific drug, all cell lines are used; cell lines that have higher matching scores with the given patient hold more weight during imputation. The weights of cell lines are determined by a flexible weight function, which is modulated by a rate parameter optimized for prediction.

**Fig. 1.**
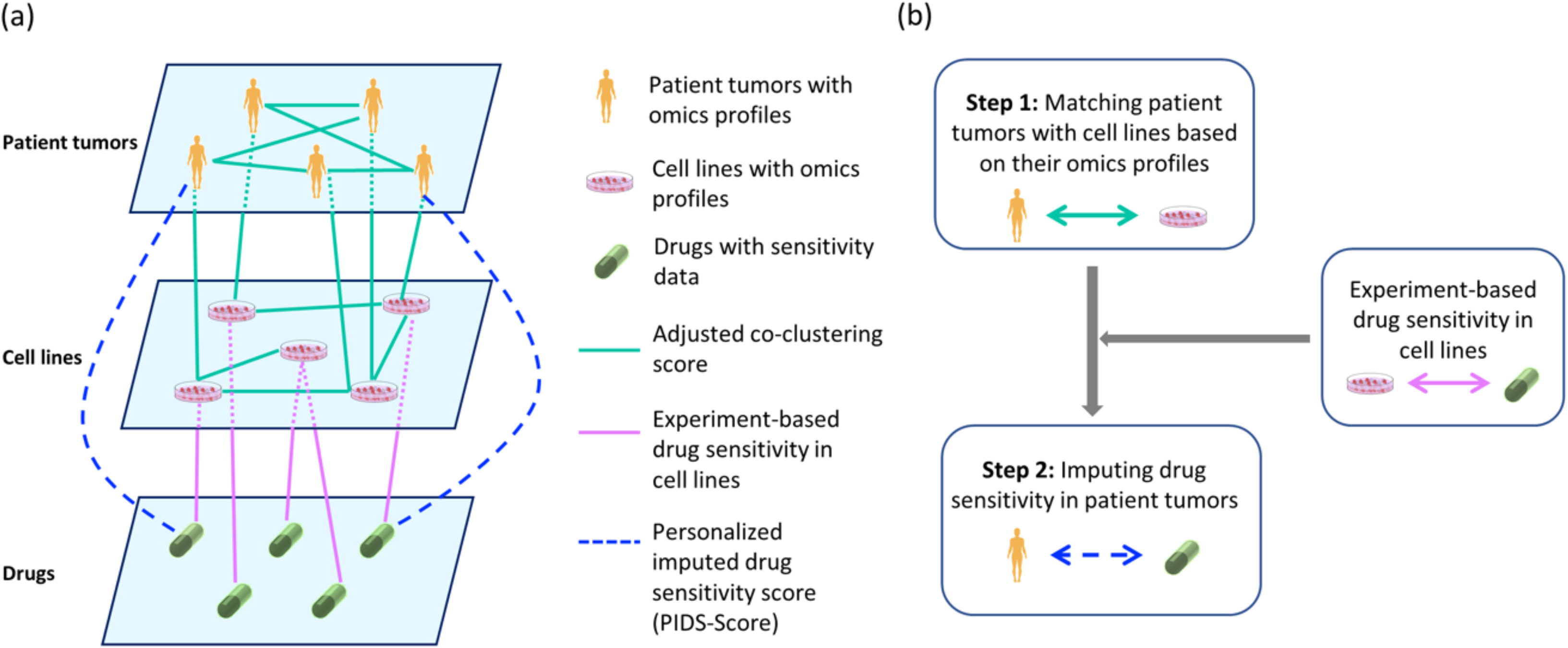
Conceptual framework (a) A conceptual organization of our multilayer networks, where the three hierarchical layers include patient tumors, cell lines and drugs (top to bottom). There are there different types of edges; edges in the network of patient tumors and cell lines (solid green lines) are estimated by our method; edges between cell lines and drugs (solid pink) are obtained from experimental data; missing edges between patients and drugs (dashed blue) are also imputed by our method. (b) Main steps of our approach. We first match patient tumors with cell lines, and then impute drug sensitivity in patient tumors based on matching results of patient tumors and cell lines as well as the dataset of drug sensitivity in cell lines.

We applied our method on 687 lung cancer patients from two subtypes of lung cancer (lung adenocarcinoma (LUAD) and lung squamous cell carcinoma (LUSC)), 648 cell lines derived from 19 different types of cancer, and 251 drugs and experimental compounds. Matching patients and cell lines can be based on any type of multi-omic data. As a proof-of-concept, we here used the Reverse Phase Protein Arrays (RPPA)-based functional proteomic data taken from The Cancer Proteome Atlas (TCPA)^12–13^ and MD Anderson Cancer Cell Lines Project (MCLP)^14^, which characterize proteins that are clinically actionable and druggable.^15^ We computed PIDS-Scores of 687 patients to all drugs. We identified the top proteomically matched cell lines and top sensitive drugs (based on their PIDS-Scores) for the entire patient cohort as well as individual patients. These identified cell lines were mainly derived from lung cancer, and the top ranked drugs were indeed highly related with the lung cancer treatment in clinics, confirming the clinical relevance of PIDS-scores. Finally, we conducted an exploratory validation by associating our PIDS-Scores with patient clinical outcomes and found significant associations between them.

## 2. Methods

### 2.1. Matching Patient Tumors with Cell Lines

In this section, we detail the steps of our analytical framework presented in Figure 2. We start with a brief overview of spectral graph wavelets. Then, we describe the algorithm of wavelet-based multiscale community detection, and matching patient tumors with cell lines based on the network-based communities. As Figure 2(a-c) shows, the omics profiles of patient tumors and cell lines (input data) are pre-processed and batch-corrected before calculating the correlations, and more details of data preprocessing and integration are deferred until Section 3.1.

**Fig. 2.**
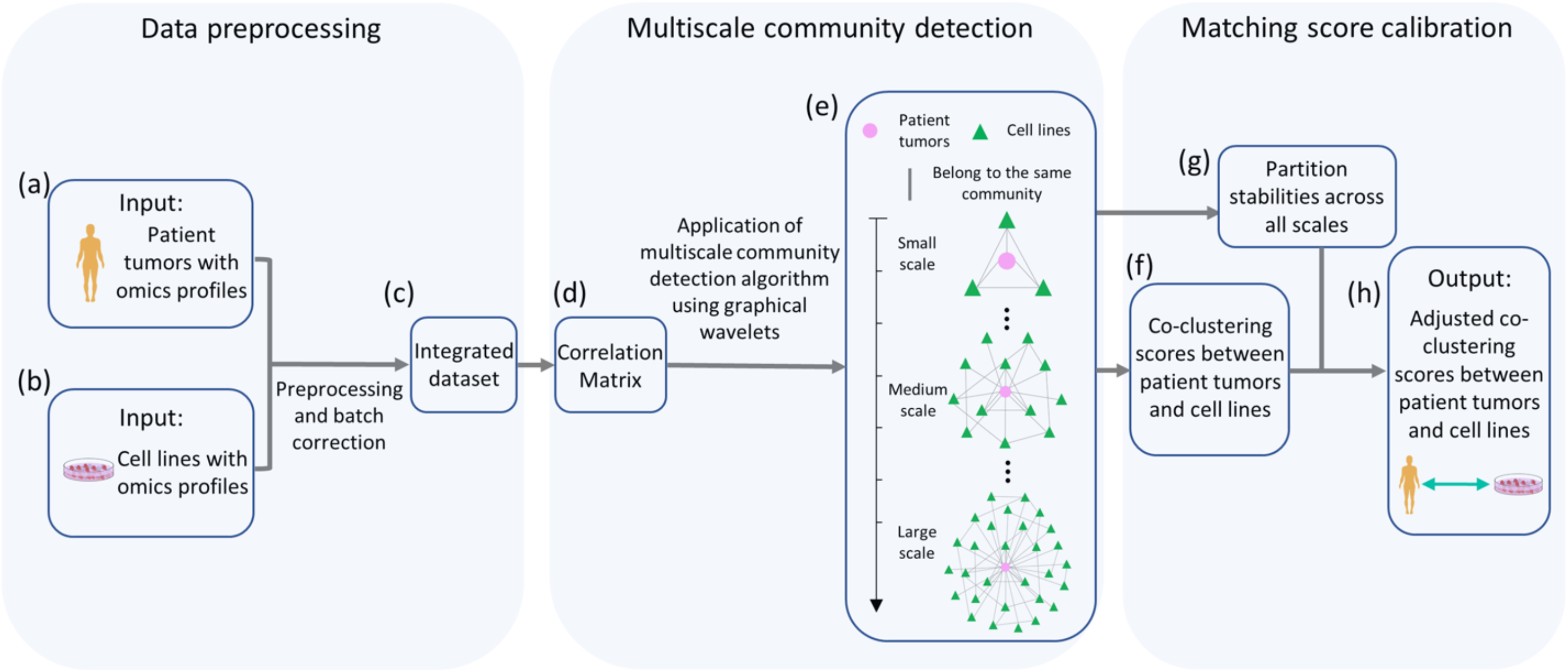
Analytical framework for matching patient tumors with cell lines. (a) – (c) Data preparation for multiscale community detection. (d) – (e) Multiscale community detection procedure. (f) – (h) Steps to calculate the adjusted co-clustering score based on the multiscale community detection results and the partition stability at each scale.

#### 2.1.1. Spectral Graph Wavelets Defined in Graph Fourier Space

Let *G* = (*V, E, A*) be an undirected, complete weighted graph with node set *V* = {*P*_1_, …, *P*_*n*_, *C*_1_, …, *C*_*m*_} (*N* = *n* + *m*), edge set *E* and weighted adjacency matrix *A*, where *P*_1_, …, *P*_*n*_ represents *n* patient tumors and *C*_1_, …, *C*_*m*_ represents *m* cell lines. To aid exposition, the indices used for *P*_1_, …, *P*_*n*_, *C*_1_, …, *C*_*m*_ are 1,2, …, *N* respectively. In this paper, we used *distance correlation*^16^ as a measure of weight (the entry of *A*) between any pair of patients and cell lines based on their omics profile (Fig. 2(d)). Unlike the Pearson correlation in which uncorrelatedness does not imply independence, a zero distance correlation implies that two random vectors are independent, and this property enables it to explain both linear and nonlinear relationships between random vectors. Additionally, distance correlation returns a bounded metric between 0 and 1. Then, the graph Laplacian matrix is defined as *L* = *D* − *A*, where *D* is the diagonal degree matrix with *D*_*ii*_ = Σ_*j*≠*i*_ *A*_*ij*_ measuring the connectivity of a certain node, and a normalized version of *L* can be constructed by taking *D*^−1/2^*LD*^−1/2^ with its eigenvalues bounded by 0 = *λ*_1_ ≤ *λ*_2_ ≤ … *λ*_*N*_ ≤ 2 and corresponding eigenvectors denoted by *χ*_1_, *χ*_2_ … *χ*_*N*_.^17^ In graph Fourier space, 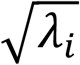 is considered as the frequency at node *i*, and *χ* = (*χ*_1_|*χ*_2_| … |*χ*_*N*_) represents the matrix of graph Fourier modes, both of which are used to leverage the multiscale information based on spectral graph wavelets.

Spectral graph wavelets were first proposed by Hammond et al.^18^, and Tremblay et al.^11^ further developed the theory and computational algorithms, which we briefly overview here. Let Ψ_*s*_ = (*ψ*_*s*,1_|*ψ*_*s*,2_| … |*ψ*_*s*,*N*_|) = χ*G*_*s*_χ^*T*^ be the wavelet basis at scale *s*, where (*ψ*_*s,a*_)_*a*=1,2,…,*N*_ represents the wavelet at scale *s* centered around node *a* or the feature vector for node *a* at scale *s*, and *G*_*s*_ = diag(*g*(s*λ*_1_), …, *g*(*sλ*_*N*_)) represents the filer based on a band-pass filter kernel *g* (⋅) and *s* from the scale set *S* = {*s*_1_, *s*_2_, …, *s*_*M*_} with scales’ total number M (M = 100 in this paper). In our analysis, we adapted the algorithm used by Tremblay et al. to determine the function *g*(⋅) as well as the scale set *S* = {*s*_1_, *s*_2_, …, *s*_1_}, and more details about the estimation can be found in their paper^11^.

#### 2.1.2. Multiscale Community Detection

Unlike spectral clustering, which only uses first k eigenvectors of *L* to construct k clusters at one scale, multiscale community detection uses different filtered information from all eigenvalues and eigenvectors of *L* at different scales based on spectral graph wavelets. At large scales, the wavelets or the node feature vectors correspond to *a coarser description of the global connectivity*; more connected components and large size communities will always be detected. Conversely, at small scales we expect small size communities, hence exploring *a more nuanced local connectivity*. Given a large network, Ψ_*s*_ computed from Section 2.1.1 is always computationally inefficient due to its high-dimensionality. Hence, we used the fast community detection procedure to obtain the set of partitions 𝒫 = {*P*_*s*_}_*s*∈*S*_ across all scales, where *P*_*s*_ represents the clustering result at scale *s*. The fast community detection algorithm, which needs to be repeated for all scales, has three steps:^11^

1. At a given scale, generate a matrix of *φ* (relatively smaller than *N*) Gaussian random vectors *r* with zero mean and variance *σ*^2^ = 1: *R* = (*r*_1_|*r*_2_| … |*r*_*φ*_ ℝ^*N*×*φ*^. Compute the dimension-reduced feature vector 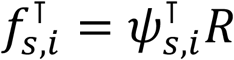 for node *i* ∈ *V*.
2. Estimate the distance matrix 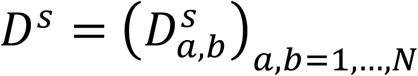 with 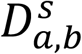 calculated by:

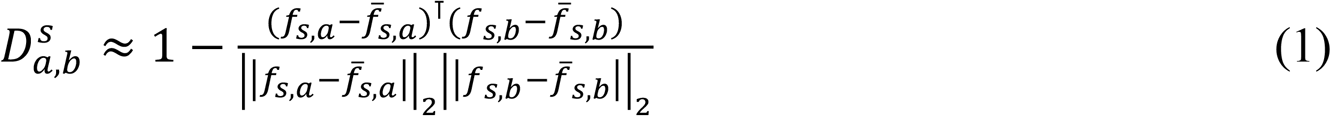

where the constant vector 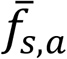 is equal to the average of *f*_*s,a*_.
3. Perform hierarchical clustering on *D*_*S*_ using a graph-cutting strategy, and obtain the set of partitions 𝒫 = {*P*_*s*_}_*s∈S*_ across all scales.

#### 2.1.3. Co-clustering Scores and Their Adjustment based on Partition Stabilities

The co-clustering score between two nodes measures the proportion of scales at which each pair of patient tumors and cell lines are clustered into the same group. Because this score combines all the network clusters at multiple scales, it is a robust measure of connectivity between nodes. Unlike the co-clustering score which treats scales the same, different weights are assigned to scales when calculating the adjusted co-clustering score; the scale, at which the graph has a higher partition stability (or clearer community structure), is given more weight. The calculation of the adjusted co-clustering score between nodes *a* and *b* is done as follows:

1. Consider *J* sets of *φ* random signals from Section 2.1.2. Compute the partition stability *γ*(*s*), which is defined as the mean similarity between all pairs of partitions of 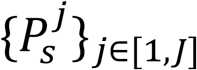.^11^
2. Repeat the step 1 for each scale *s*. The adjusted co-clustering score for nodes a and b is:

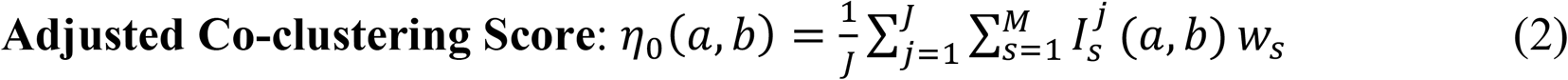

where 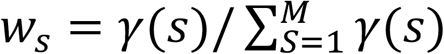 is the weight given to the scale *s*, and 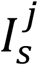 is the indicator function. 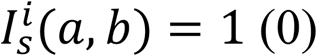 if nodes *a* and *b* are in the same (different) cluster (s) at scale *s* of partition *j*.
3. Get *η*_0_ for all pairs of nodes, and then multiply the largest *η*_0_ to 1 by a constant number *k*. The final adjusted co-clustering score for nodes a and b is *η*(*a*, *b*) = *kη*_0_(*a*, *b*). The reasoning for this adjustment is that the only meaningful metric is the relative difference between patient-cell line co-clustering levels.

Since patients and cell lines serve as nodes in our network, the adjusted co-clustering scores are computed for all pairs based on the algorithm in Section 2.1.

### 2.2. Imputing Drug Sensitivity in Individual Patients

This section details the steps of imputing patient drug sensitivity (Figure 3), where the co-clustering results from Section 2.1 serve as our input dataset. For clarity of exposition, we only introduce the pipeline of imputing the sensitivity of a given drug *D* in an individual patient *P*. As Figure 3(a) shows, our imputation method takes advantage of network topologies (1) between patient *P* and cell lines, (2) within cell lines, and (3) between cell lines and drug *D*.

**Fig. 3.**
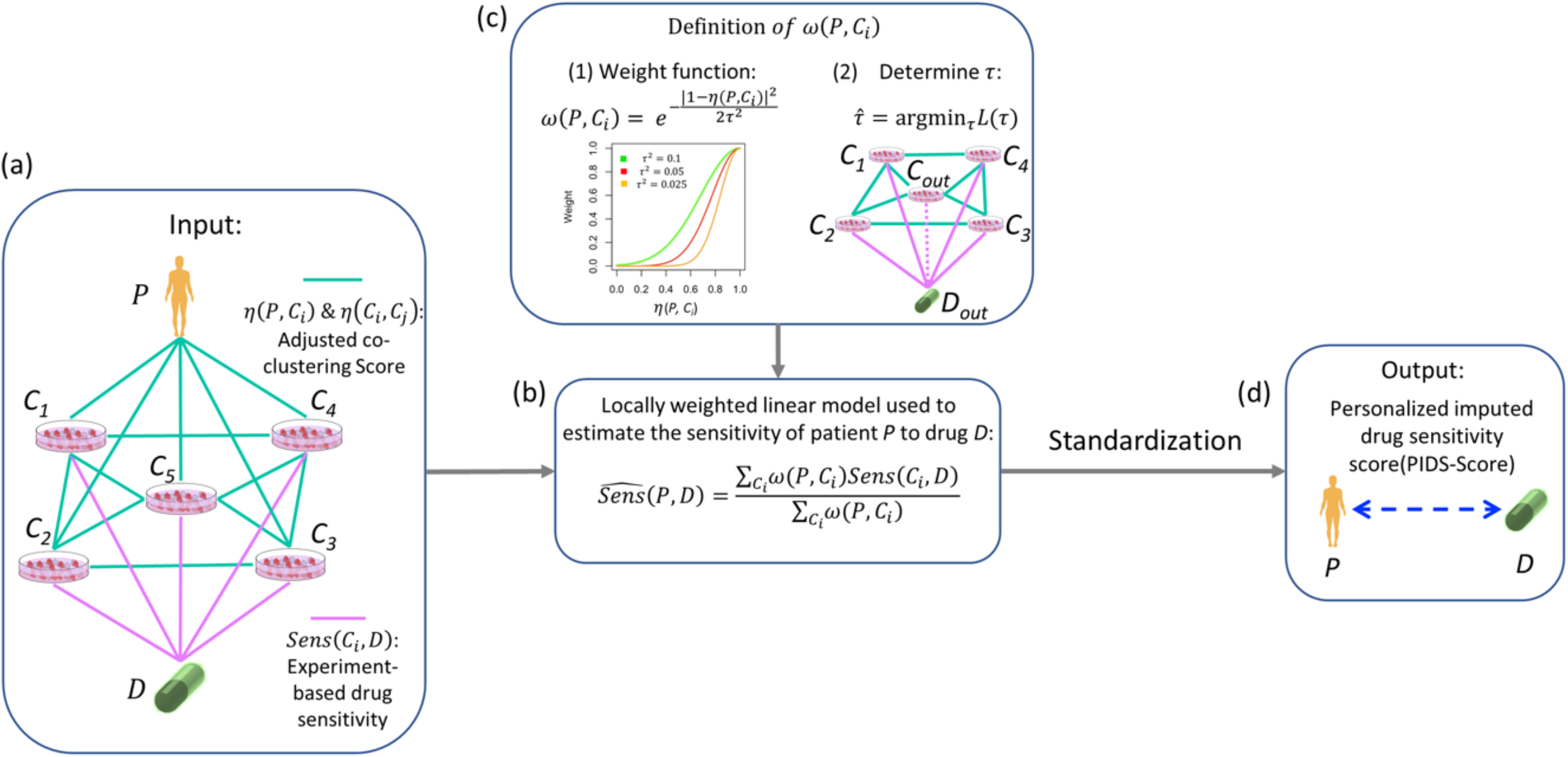
PIDS-Score computation for a given patient-drug pair. (a) Input datasets for the computation include the patient-cell line matching results and the experiment-based sensitivities of drug *D* in cell lines. (b) – (c) A locally weighted linear model is used to estimate the sensitivity of drug *D* for patient *P*. The weight between patient *P* and cell line *C*_*i*_ in the model decreases as the matching score *η*(*P*, *C*_*i*_) decreases, as shown in the left sub-panel (1). The right sub-panel (2) shows that the rate parameter *τ* in the weight function is derived from an objective function using leave-one-out cross-validation, where the pink dashed line is the link to be predicted in each iteration. (d) Finally, the result from the locally weighted linear model is standardized to obtain the PIDS-Score that is used for interpretation and inference.

#### 2.2.1. Locally Weighted Linear Model

Recent work has utilized a locally weighted linear model based on a dual-layer integrated cell line-drug network to predict the drug sensitivity of a new cell line to a known drug.^19^ In this study, we generalized this model to a new locally weighted linear model based on our three-layer network so that the drug sensitivity of patient *P* to drug *D* can be imputed. Cell lines to be used are those available in both patient *P*-cell line network and cell line-drug *D* network (Figure 3(a)). Under the assumption that a given patient tumor and its highly matching cell lines will react similarly to the same drug, the main formula of the locally weighted linear model used to impute the drug sensitivity of patient *P* to drug *D* is:

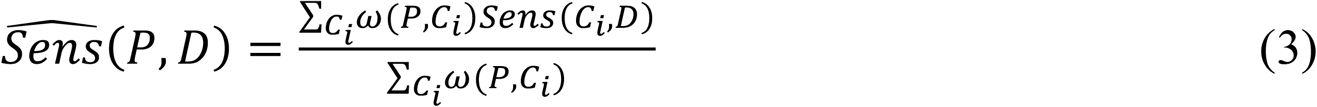

where *ω*(*P*, *C*_*i*_) is the weight parameter based on the adjusted co-clustering score between patient *P* and cell line *C*_*i*_, and *Sens*(*C*_*i*_, *D*) is the experiment-based sensitivity of cell line *C*_*i*_ to drug *D* for which any appropriate drug sensitivity metrics can be used.^19^ Since there is no agreement with respect to the optimal metric for summarizing the information from the dose response curve, in this study we used pIC_50_ = −logIC_50_, a commonly used metric to measure the potency of compound inhibition.^20^ Specifically, IC_50_ represents the concentration of a drug required for 50% inhibition of cell growth, and a larger pIC_50_ corresponds to a more potent compound.

In order to build an appropriate and flexible weight function of the adjusted co-clustering score between patient *P* and cell line *C*_*i*_, we followed two criterions: 1) the function should be increasing since a cell line *C*_*i*_ highly matched to patient *P* is expected to have a high impact on the imputation of drug sensitivity in patient *P*; 2) the function should be flexible according to nonlinear dependency controlled by one or more free parameters. Similar to the “weight” in locally weighted linear regression, an appropriate choice of the weight function in our model is 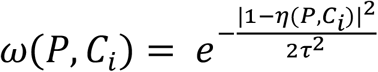, where *η*(*P*, *C*_*i*_) is the adjusted co-clustering score between patient *P* and cell line *C*_*i*_, and *τ* is the rate parameter which controls the rate at which *ω*(*P*, *C*_*i*_) decays with distance of patient *P* from cell line *C*_*i*_.^19^ As shown in Figure 3(c), if *η*(*P*, *C*_*i*_) is close to 1, *ω*(*P*, *C*_*i*_) will be close to 1, which means the cell line *C*_*i*_ has a great contribution to imputing the drug sensitivity in patient *P*. In contrast, if *η*(*P*, *C*_*i*_) is small, *ω*(*P*, *C*_*i*_) could be close to 0 even though *η*(*P*, *C*_*i*_) is not close to 0, which implies that a cell line moderately matched with patient *P* may have very low impact on the imputation.

##### Rate parameter determination

The rate parameter *τ* in the weight function *ω* is determined by leave-one-out cross-validation, which requires both observed and predicted drug responses. Since true patient drug sensitivity data are not available in our model, only the network of cell lines and drugs is used to find the best value of *τ*. However, because patient tumors and cell lines have the same type of omics profile and the same definition of edges in their network, we considered the value of *τ* determined by the cell line-drug network as an appropriate choice of *τ* used to impute the drug sensitivity of patient *P* to drug *D*. In the leave-one-out cross-validation procedure^19^, each cell line-drug pair is held out to serve as the test data while other edges in the cell line-drug network are used to predict the link between the cell line and the drug that are left out.

#### 2.2.2. Standardization

For different drugs, their values of pIC_50_ to a given set of cell lines are potentially on different scales, which percolates to the imputed patient drug sensitivity 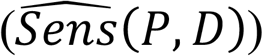 derived from these values (Eq. (3)). To standardize 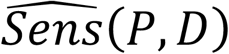, it is then compared to the sensitivities of drug *D* in all cell lines. Similar to the formula for computing z-scores, the standard function is defined as follows:

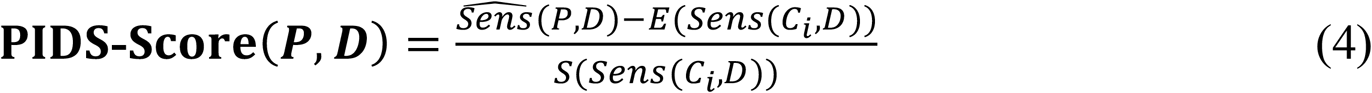

where 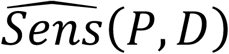 is the (unstandardized) imputed drug sensitivity, *E*(*Sens*(*C*_*i*_, *D*) is the mean value of the sensitivities of drug *D*, and *S*(*Sens*(*C*_*i*_, *D*)) is the standard deviation of the sensitivities of drug *D* across all cell lines. The PIDS-Score measures how many standard deviations from the imputed drug sensitivity of drug *D* in patient *P* to the mean sensitivity. Similar to z-scores, PIDS-Scores always fall between −2 and 2 where a higher value indicates a higher drug sensitivity.

## 3. Results

### 3.1. Functional Proteomics and Drug Sensitivity Datasets

In our approach, the two main steps (Figure 1(b)) use different types of input datasets. First, while our method of matching patient tumors with cell lines can be applied on any types of omics profiles, here we used Reverse Phase Protein Array (RPPA)-based functional proteomics data for two primary reasons: 1) this type of data can adequately capture downstream aberrations of proteins missed by (upstream) genomics and transcriptomics data; 2) aberrations in proteomic markers are more closely related to eventual clinical phenotypes or outcomes. The dataset of patient tumors taken from TCPA^12,13^ contains >8,000 samples of 32 cancer types, and the dataset of cell lines taken from MCLP^14^ includes >650 independent cell lines derived from 19 different tissues. To illustrate our approach, we focused on two subtypes of lung cancer: lung adenocarcinoma (LUAD) and lung squamous cell carcinoma (LUSC), both of which are non-small cell lung cancer (NSCLC), one of the most common human cancer types. The combined proteomics datasets from TCPA and MCLP were then pre-processed for missing value imputation and batch correction using kNN imputation^21^ and ComBat^22^, respectively. After preprocessing, there are 687 patient tumors, including 362 LUAD samples and 325 LUSC samples; 648 cell lines derived from 19 different tissues; and 165 overlapping proteins, which serve as features for patient tumors and cell lines. In addition, the drug sensitivity data for cell lines were obtained from GDSC portal, which provides the drug sensitivities (IC_50_) of 251 drugs and compounds in 293 cell lines recorded by both MCLP and GDSC.

### 3.2. Network-based Matching of LUAD and LUSC Patients with Cell Lines

Following the pipeline in Section 2.1, we matched 687 NSCLC patients with 648 cell lines using their functional proteomics profiles, and obtained adjusted co-clustering scores between all patient-cell line pairs. Figure 4(a) shows a heat map of the adjusted co-clustering scores (between 0 to 1), where rows represent patients from the two subtypes (shown in green and purple bands on the left) and columns represent cell lines derived from 19 different tissues. We observed a high-level connectivity (colored in red) between LUAD patients and two types of cell lines, *lung* and *head and neck*, which is consistent with previous findings^1^ that head and neck cancer samples and many lung cancer samples belong to the same molecular subgroup. In contrast, adjusted co-clustering scores are only high between LUSC patients and a few cell lines mainly derived from lung tumors.

**Fig. 4.**
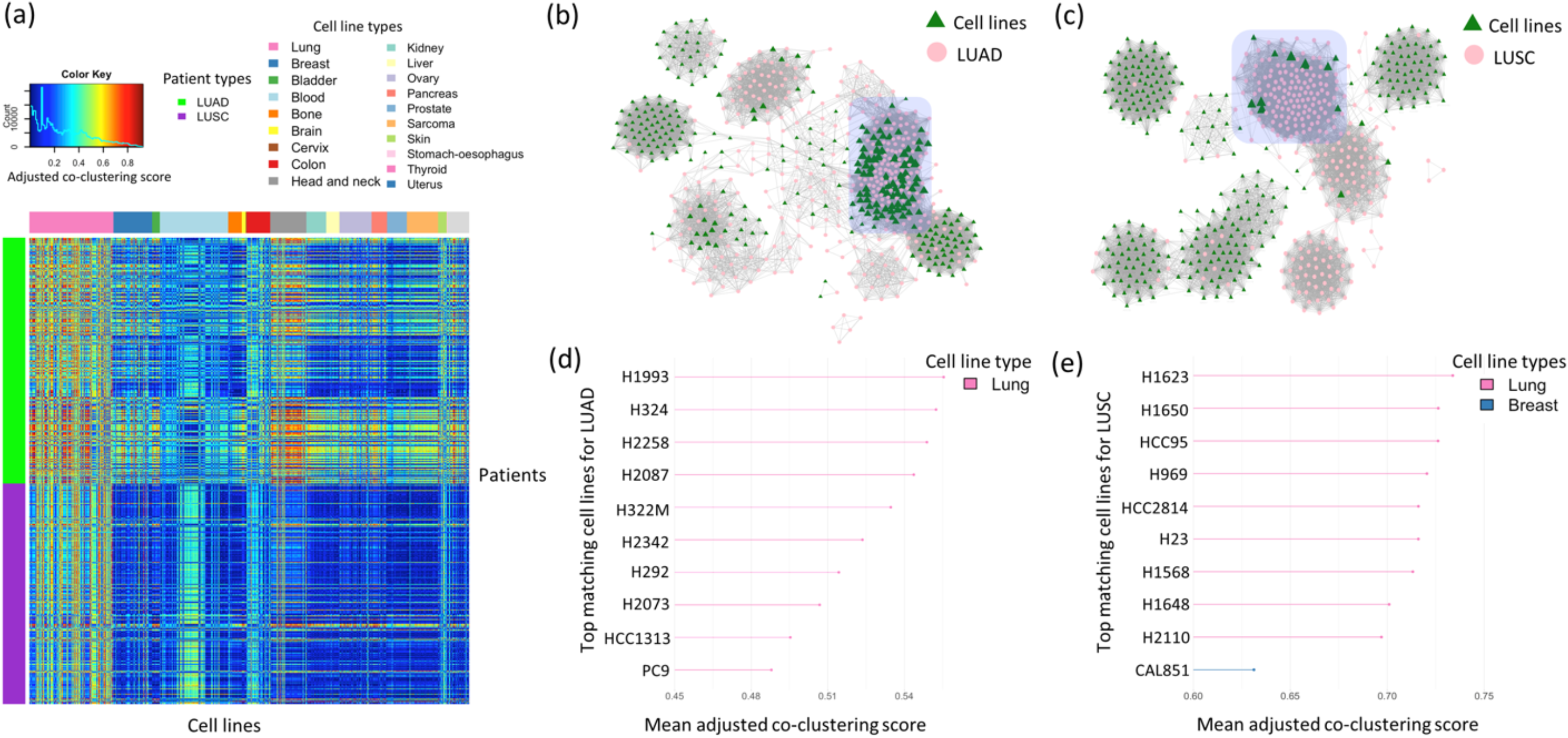
Network-based matching of patients and cell lines. (a) Heat map showing the adjusted co-clustering scores between patient tumors and cell lines for both LUAD and LUSC subtypes. (b) – (c) LUAD patient-cell line network and LUSC patient-cell line network. (d) – (e) Bar plots showing the top matching cell lines for LUAD and LUSC patient samples. A mean adjusted co-clustering score is equal to the average of adjusted co-clustering scores between a given cell line and all patient tumors in a given subtype.

We also plotted the patient-cell line networks for the two subtypes of lung cancer. As shown in Figure 4(b-c), the edge between two samples is connected when their adjusted co-clustering score is higher than 95% of scores computed from all pairs of samples; the size of a green triangle (cell line) depends on the number of edges connected to it. In the LUAD patient-cell line network (Figure 4(b)), there is a relativity large community (highlighted) consisting of approximately 100 patient samples and approximately 70 cell lines. In Figure 4(c), the LUSC patient-cell line network has a clearer community structure than LUAD with 6-8 well separated communities, where one of the communities (highlighted) has relatively large size triangles (cell lines connected to >80 LUSC patients). From these networks, we selected the top 10 matching cell lines for the LUAD and LUSC cohorts, based on the mean adjusted co-clustering scores (Figure 4(d-e)).

For LUAD, unsurprisingly, all listed cell lines are derived from lung tumors (Figure 4(d)). H1993 (ranked #1 based on mean adjusted co-clustering scores), a cell line derived from a metastatic LUAD patient, has been frequently used to study MET gene activation in NSCLC.^23,24^ For LUSC (Figure 4(e)), while most of the top matching cell lines were derived from lung tumors, one highly matching cell line was derived from a breast tumor (CAL851, ranked #10). H1623, the top matching cell line for LUSC patients, was derived from a metastatic NSCLC patient, and H1650 (ranked #2) has often been used to test the efficacy of gefitinib in inhibiting EGFR pathway.^23,25^

### 3.3. Drug Sensitivity Imputation for LUAD and LUSC Patients

To impute the drug sensitivity scores for patients (Section 2.2), we selected 293 overlapping cell lines with drug response profiles across MCLP and GDSC (as training data) and used the patient-cell line matching results (above) to compute PIDS-Scores of the 362 LUAD patients and 325 LUSC patients to 251 different drugs and experimental compounds. In Figure 5(a), we plotted the heat map of PIDS-Scores to provide a global overview of the imputed drug responses across the 251 drugs and compounds for both cohorts. As Figure 5(a) shows, there are three clear patterns: 1) for a given drug, its PIDS-Scores appear to be consistent across patients from the same cohort – clear vertical (column-wise) banded structure in the heat map; 2) among different drugs, their PIDS-Scores vary significantly for a given patient (row-wise); and 3) within each subtype of patients, a few (2-3) patient sub-clusters show different patterns of overall drug response profiles, indicating the existence of within-subtype heterogeneity with regard to drug responses.

**Fig. 5.**
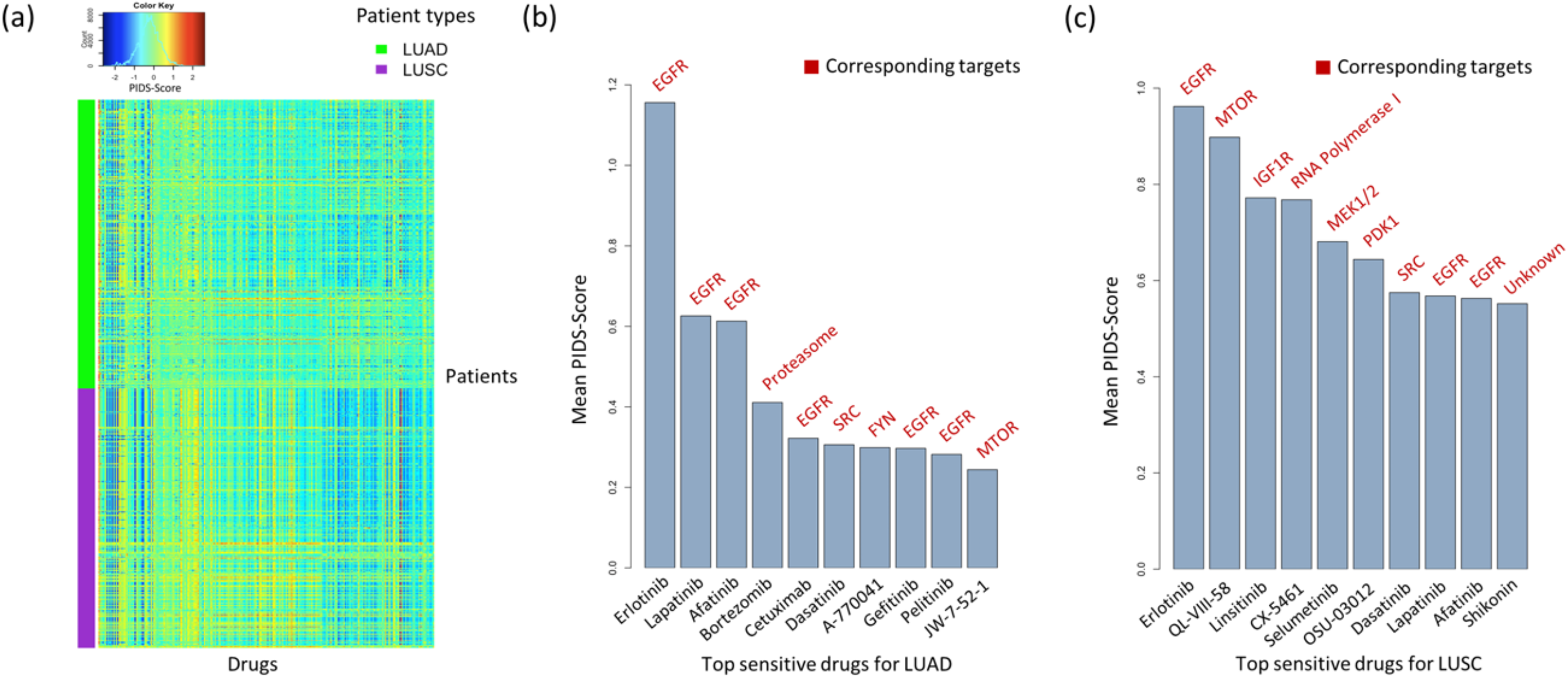
Visualization and summary of PIDS-Scores. (a) Heat map showing PIDS-Scores of all patient-drug combinations. (b) – (c) Bar plots showing the top sensitive drugs for LUAD and LUSC patient samples based on the mean PIDS-Scores. A mean PIDS-Score is equal to the average of PIDS-Scores between a given drug and all patient tumors in a given subtype. For listed drugs, their corresponding targets are also listed above the bars.

Based on the mean PIDS-Scores, we selected top 10 sensitive drugs for both LUAD and LUSC patients, which are shown in Figures 5(b) and 5(c) respectively. A total of 16 unique drugs are ranked in these two top 10 lists. Three of them (*erlotinib, afatinib and gefitinib*) have been approved by the Food and Drug Administration (FDA) for the treatment of NSCLC.^26^ In addition, multiple clinical and translational studies have verified that all listed targets of these 16 drugs are related with the pathogenesis of lung cancer; EGFR, which appears 9 times as various top drugs’ targets in Figure 5(b-c), is frequently overexpressed in lung cancer, and many NSCLC studies over the last decade have been focused on treating EGFR mutations;^27^ proteasome, a protein degradation complex, has become an attractive therapeutic target in both NSCLC and small cell lung cancer (SCLC);^28^ mTOR is a kinase which regulates normal cell growth, and its dysregulation often occurs in lung cancer;^29^ IGF1R, a type of tyrosine kinase receptors, is also highly expressed in lung cancer.^30^

#### Patient-specific analyses

The PIDS-Scores can also be used for patient-specific inferences and potential drug recommendations, above and beyond the assessment of average drug sensitivity across patient cohorts. To facilitate this functionality, we have developed a web-based visualization tool using R-shiny (https://qingzliu.shinyapps.io/psb-app/) to provide patient-specific drug summaries. In Figure 6, we plotted two flow diagrams to show one patient example, an LUAD patient (TCGA ID: TCGA-44-2657). The left panel demonstrates the connectivity between a given patient, the patient’s top matching cell lines and PIDS-Score based top sensitive drugs. Among the top selected drugs for this patient, three of them (*vinorelbine, trametinib and docetaxel*) have been approved by FDA for NSCLC treatments.^26^ The target of *trametinib* is MEK1/2 while the target of *vinorelbin*e and *docetaxel* is microtubule stabilizer. Interestingly, both of these targets are not among the listed top 10 drugs for LUAD patients shown in Figure 5(b). This highlights the fact that even within a given clinical subtype, patients could be heterogenous and thus respond differently to different drugs – hence emphasizing the importance of precision medicine-driven strategies.^31^

**Fig. 6.**
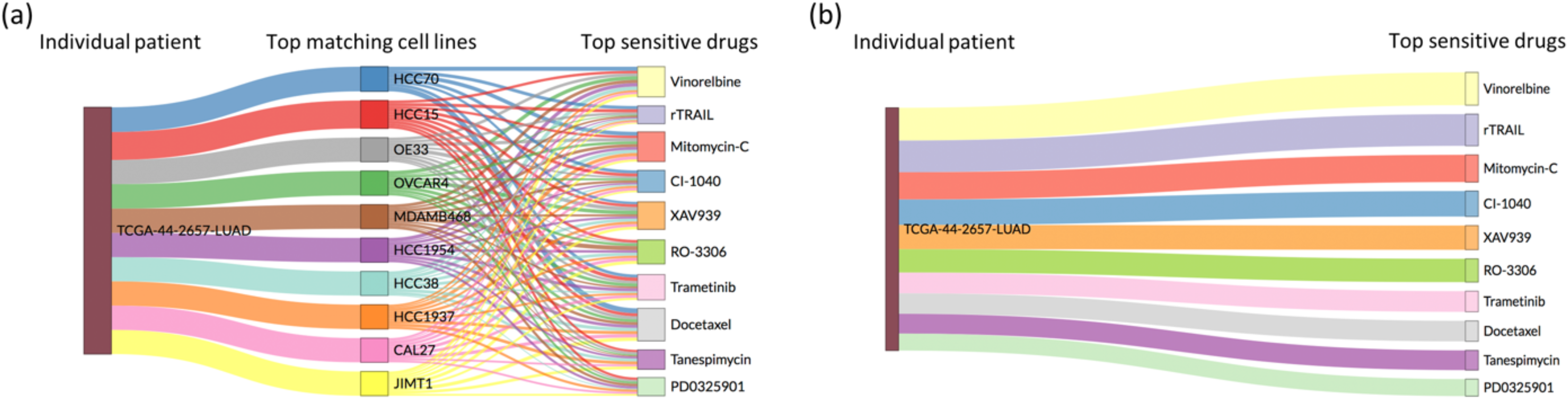
Flow diagrams illustrating the relationship between a given patient, the patient’s top matching cell lines and top sensitive drugs. The widths of edges between the patient and cell lines are based on the adjusted co-clustering scores (Panel a); the widths of edges between the patient and drugs are based on the PIDS-Scores (Panel b).

### 3.4. Association with Ex-vivo Patient Outcomes

Given the lack of *in-vivo* drug response data from patients, there was no absolute ground truth for validating our PIDS-Scores. As an exploratory validation, we associated PIDS-Scores of each drug across all LUAD or LUSC patients with their survival outcomes (survival times and statuses), using univariate Cox proportional hazards models. We found that the PIDS-Scores of LUSC patients to *alectinib* (*β* = −1.50, P-value < 0.001), *CX-5461* (*β* = −0.63, P-value < 0.001), and *linsitinib* (*β* = −0.61, P-value = 0.02) are significantly associated with their survival data. These regression coefficients are negative and hence in the right direction, i.e. higher (lower) PIDS-Scores are associated with longer (shorter) survival times, indicating their relative therapeutic potential. Indeed, *alectinib* has been approved by FDA for the treatment of NSCLC.^26^

## 4. Discussion

We propose a network-based approach to identify, calibrate and narrow the therapeutic potential of investigational drugs, by integrating omics data across patient tumors and cancer cell lines. Using a robust multi-scale strategy, we identified best representative cell lines and used this information to impute *in-vivo* drug sensitivities (PIDS-Scores) for individual patients from two subtypes of lung cancer. We show that our identified cell lines and top sensitive drugs are in concordance with established lung cancer cell lines and drugs with lung cancer-related targets. We also found significant associations between PIDS-Scores and patient clinical outcomes. These results suggest that our new computational approach is promising to help optimize the individual patient-drug recommendations and to aid precision oncology endeavors.

Our approach can be generalized and extended to other model systems and pan-omic data. For example, currently the imputation is based on cell lines, which might not entirely recapitulate the complex microenvironment in patient tumors.^8^ More clinically relevant systems such as organoids and patient derived xenografts, which can better recapitulate human physiology, could be used as these datasets mature. We also plan to expand our analyses to other cancer types and molecular levels, and investigate a wider range of drugs and experimental compounds. In addition, we can incorporate drug-drug interactions in the bottom layer of our network, which help further identify omics-guided combinational therapies at an individual patient level.

## Acknowledgments

The authors acknowledge and thank Neel Desai and Dr. Jeffrey S. Morris for their inputs on the graphical wavelet models and Drs. Rehan Akbani, Han Liang and Gordon B. Mills for giving us access to the RPPA datasets.

## References

1. K. A. Hoadley, C. Yau, et al., Cell 158, 929 (2014).

2. J. Yang, A. Li, Y. Li, X. Guo, M. Wang, Bioinformatics 35, 1527 (2019).

3. The Cancer Genome Atlas homepage; http://cancergenome.nih.gov/abouttcga

4. The International Cancer Genome Consortium, Nature 464, 993 (2010).

5. J. C. Costello, L.M. Heiser, et al., Nature Biotechnology 32, 1202 (2014).

6. J. Barretina, G. Caponigro, et al., Nature 483, 603 (2012).

7. W. Yang, J. Soares, et al., Nucleic Acids Research 41, D955 (2013).

8. A. Goodspeed, L. M. Heiser, et al., Molecular Cancer Research 14, 3 (2016).

9. F. Lorio, U. McDermott, et al., Cell 166, 740 (2016).

10. P. Geeleher, N. J. Cox, R. S. Huang, Genome Biology 15, R47 (2014).

11. N. Tremblay, P. Borgnat, IEEE Transactions on Signal Processing 62, 5227 (2014).

12. J. Li, R. Akbani, W. Zhao, Y. Lu, J. Weinstein, et al., Cancer Research 77, e51 (2017).

13. J. Li, Y. Lu, et al., Nature Methods 10, 1046 (2013).

14. J. Li, W. Zhao, et al., Cancer Cell 31, 225 (2017).

15. H. Masuda, Y. Qi, et al., Oncotarget 8, 70481 (2017).

16. G. J. Székely, M. L. Rizzo, N. K. Bakirov, The Annals of Statistics 35, 2769 (2007).

17. F. R. Chuang, American Mathematical Society 92 (1994).

18. D. Hammond, et al., Applied and Computational Harmonic Analysis 30, 129 (2011).

19. N. Zhang, H. Wang, et al., PLOS Computational Biology 11, e1004498 (2015).

20. M. Fallahi-Sichani, S. Honarnejad, et al., Nature Chemical Biology 9, 708 (2013).

21. T. Hastie, et al., Stanford University Statistics Department Technical Report (1999).

22. W. E. Johnson, A. Rabinovic, C. Li, Biostatistics 8, 118 (2007).

23. R. M. Phelps, B.E. Johnson, et al., Journal of Cellular Biochemistry. Supplement 24, 32 (1996).

24. B. Lutterbach, Q. Zeng, et al., Cancer Research 67, 2081 (2007).

25. Y. J. Choi, J. K. Rho, et al., Cancer Chemotherapy and Pharmacology 66, 381 (2010).

26. The Food and Drug Administration homepage; https://www.fda.gov

27. B. A. Chan, B. G. Hughes, Translational Lung Cancer Research 4, 36 (2015).

28. G. Scagliotti, Critical Reviews in Oncology/Hematology 58, 177 (2006).

29. S. Ekman, M. W. Wynes, F. R. Hirsch, Journal of Thoracic Oncology 7, 947 (2012).

30. F. Nurwidya, S. Andarini, et al., Malaysian Journal of Medical Sciences 23, 9 (2016).

31. Q. Ma, A. Y. Lu, Pharmacological Reviews 63, 437 (2011).

